# Compartmentalized glycogenolysis regulates lung cancer transcription

**DOI:** 10.1101/681833

**Authors:** Ramon C. Sun, Vikas V. Dukhande, Shane Emanuelle, Lyndsay E. Young, Christine Fillmore-Brainson, Matthew S. Gentry

## Abstract

The role of cellular metabolites in the direct control of signaling is an emerging and rapidly evolving field. Herein, we identify a key role for nuclear glycogen in epigenetic regulation through compartmentalized acetyl CoA production and histone acetylation. Nuclear glycogenolysis is dependent on ubiquitination and translocation of glycogen phosphorylase (GP) into the nucleus by malin, an E3 ubiquitin ligase. We developed an innovative *in organello* stable isotope tracer method coupled to mass spectrometry analysis to define the metabolic fate of nuclear glycogen. This work revealed that GP is required for nuclear glycogen degradation and subsequent glycolysis to generate substrates for histone acetylation. Inhibition of nuclear glycogenolysis is found to be particularly important in non-small cell lung cancer (NSCLC), as evident by increased nuclear glycogen accumulation and malin suppression in NSCLC. Re-introduction of malin in model NSCLC cell lines restores nuclear glycogenolysis, resulting in increased histone acetylation and transcriptional changes that delay cancer cell growth *in vivo*. This study uncovers a previously unknown role for glycogen metabolism in the nucleus and elucidates another way by which cellular metabolites control epigenetic regulation.

---

Glycogen is the primary source of storage carbohydrate in mammals and primarily functions as an energy cache. Abnormal glycogen metabolism has deleterious effects in a range of diseases including cancer^1,2^, neurodegeneration^3^ and congestive heart failure^4^. However, glycogen also has other key roles beyond functioning as a simple energy reserve. Nuclear glycogen was first reported in the 1950s in hepatocytes, suggesting compartmentalized regulation, but its role in cellular physiology remained unanswered^5,6^. This study investigates the half-century old phenomenon of nuclear glycogen in the case of non-small cell lung cancer (NSCLC). We found increased glycogen accumulation in the nucleus of NSCLC patient tumors and model cell lines and defined its pivotal role in modulating histone acetylation.

We analyzed paired cancer and normal distal benign tissues from four NSCLC patients and three NSCLC cell lines for glycogen content. A general increase in glycogen is observed in cancer tissue and cell lines by histochemical imaging with an anti-glycogen antibody and biochemical quantification (supplement Fig. 1 and 2A-B). Further analysis of histochemical staining suggested the presence of nuclear glycogen in NSCLC (supplement Fig. 2A). To quantify the presence of nuclear glycogen, we used a hypotonic cell lysis buffer to isolate nuclear glycogen and measured the glycogen content in both cancer and normal tissues and lung cancer cell lines. Nuclear glycogen content ranged from 1-10mg/mg protein in both patient cancer tissues and cancer cell lines (Fig. 1A and D), representing a 10-100-fold increase compared with benign tissue.

**Figure 1:**
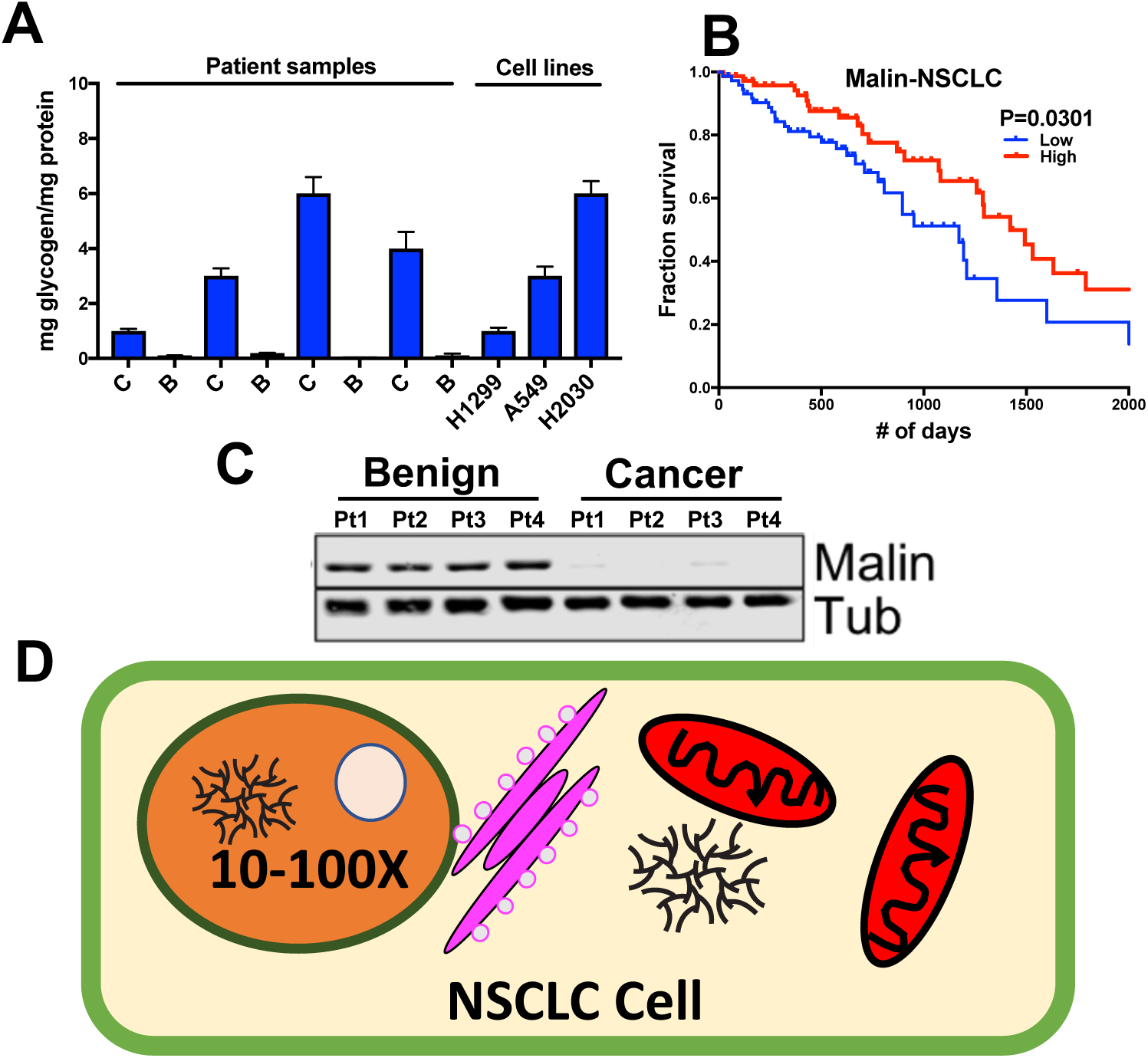
(**A**) Nuclear glycogen content in 4 pairs of NSCLC patient samples (C: cancer, B: benign) and 3 lung cancer cell lines. (**B**) Kaplan and Meier analysis of malin expression versus overall survival of NSCLC patients. (**C**) Malin expression in the same 4 pairs of NSCLC patient samples as in A. (**D**) Diagram representation of the 10-100-fold increase in compartmentalized glycogen in NSCLC.

**Figure 2:**
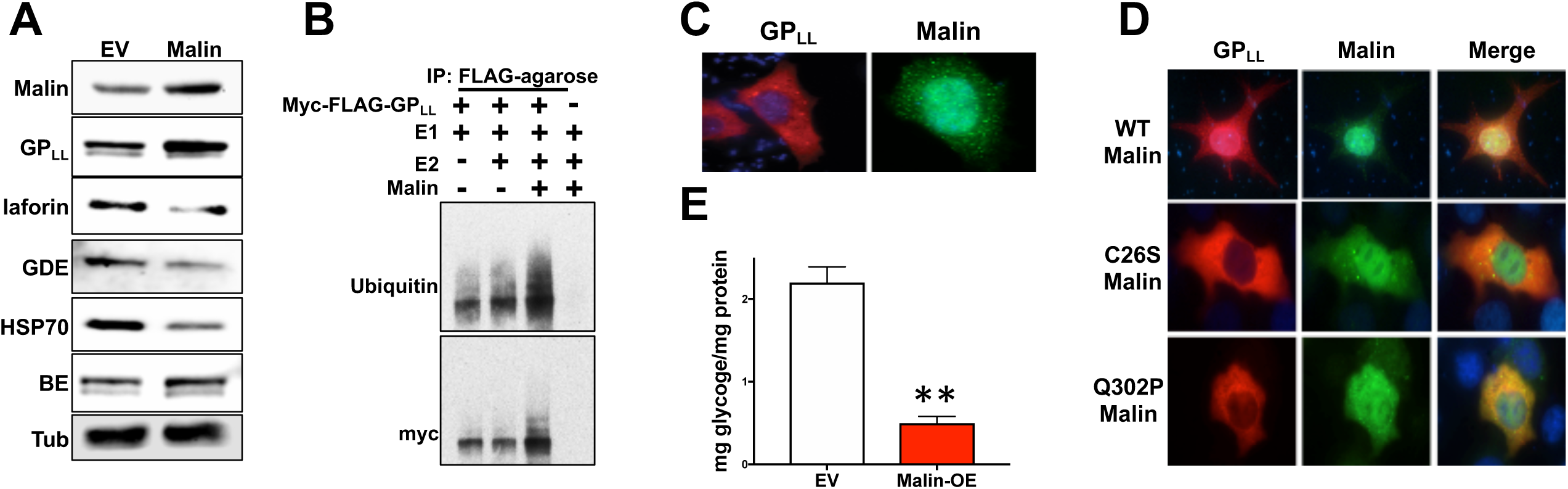
(**A**) Immunoblotting analysis of HSP70, laforin, glycogen debranching enzyme (GDE), branching enzyme (BE) and glycogen phosphorylase liver isoform (GP_LL_) with *α*-tubulin (Tub) as the loading control in cells with empty vector (EV) or malin overexpression (Malin). (**B**) *In vitro* ubiquitination assay using recombinant E1, E2 (UbcH5a), malin, and myc-GP_LL_ followed by immunoprecipitation with anti-myc, SDS-PAGE, and immunoblotting analysis with anti-ubiquitin and anti-myc antibodies. (**C**) Localization of exogenously expressed malin and GP_LL_ in HEK293 cells assessed by immunofluorescence. (**D**) Malin promotes nuclear localization of GP_LL_. GP_LL_ was co-expressed with either malin wildtype (WT) or malin mutants, C26S or Q302P, in HEK293 cells and protein localization was assessed by immunofluorescence. (**E**) Quantification of nuclear glycogen in the absence (EV) and presence of exogenously expressed malin (malin-OE). ^*^0.01 < *P* < 0.05; ^**^ 0.001 < *P* < 0.01; ^***^ *P* < 0.001, two-tailed *t*-test (see Methods)

Nuclear glycogen accumulation is accompanied by the suppression of malin, an E3 ubiquitin ligase reported to interact with glycogen metabolic enzymes^7,8^. Using OncoLnc^9^, which connects TCGA survival data with matching RNAseq analysis, we found that high malin mRNA expression correlates with better survival, suggesting malin is of clinical importance in NSCLC (Fig. 1B). Strikingly, this correlation was specific, since it was not observed with other cancers, such as breast (supplement Fig. 2D), prostate or ovarian cancers. Indeed, we observed a dramatic decrease in malin protein levels in NSCLC patient samples by both immunoblotting (Fig. 1C), and histochemical staining (supplement Fig. 1B).

Ubiquitination can impact protein levels by promoting proteasome-dependent degradation, can change enzymatic activity, and/or can change the localization of a protein^10^. Therefore, we hypothesized that downregulation of malin contributes to nuclear glycogen accumulation by impacting one or more glycogen metabolic enzymes via at least one of these mechanisms. Malin is comprised of a RING domain, a linker region, and six NHL protein-protein interaction repeats (supplement Fig. 3A)^7^. To identify novel malin substrates that could modulate nuclear glycogen, we purified recombinant malin, incubated the protein with cell extract, and identified malin-bound proteins by mass spectrometry. The majority of identified interacting proteins were previously reported including the glycogen phosphatase laforin, glycogen debranching enzyme (GDE), and HSP70. Glycogen phosphorylase (GP) and glycogen branching enzyme (GBE) were identified as novel putative malin-interacting partners (supplement Fig. 3A and B). Using co-expression and co-immunoprecipitation experiments, we validated the known interactions and the malin interaction with GP^8^. Malin over-expression (OE) resulted in decreased levels of laforin, HSP70, and GDE in our model cell line (Fig. 2A), validating previous reports^7,8^.

**Figure 3:**
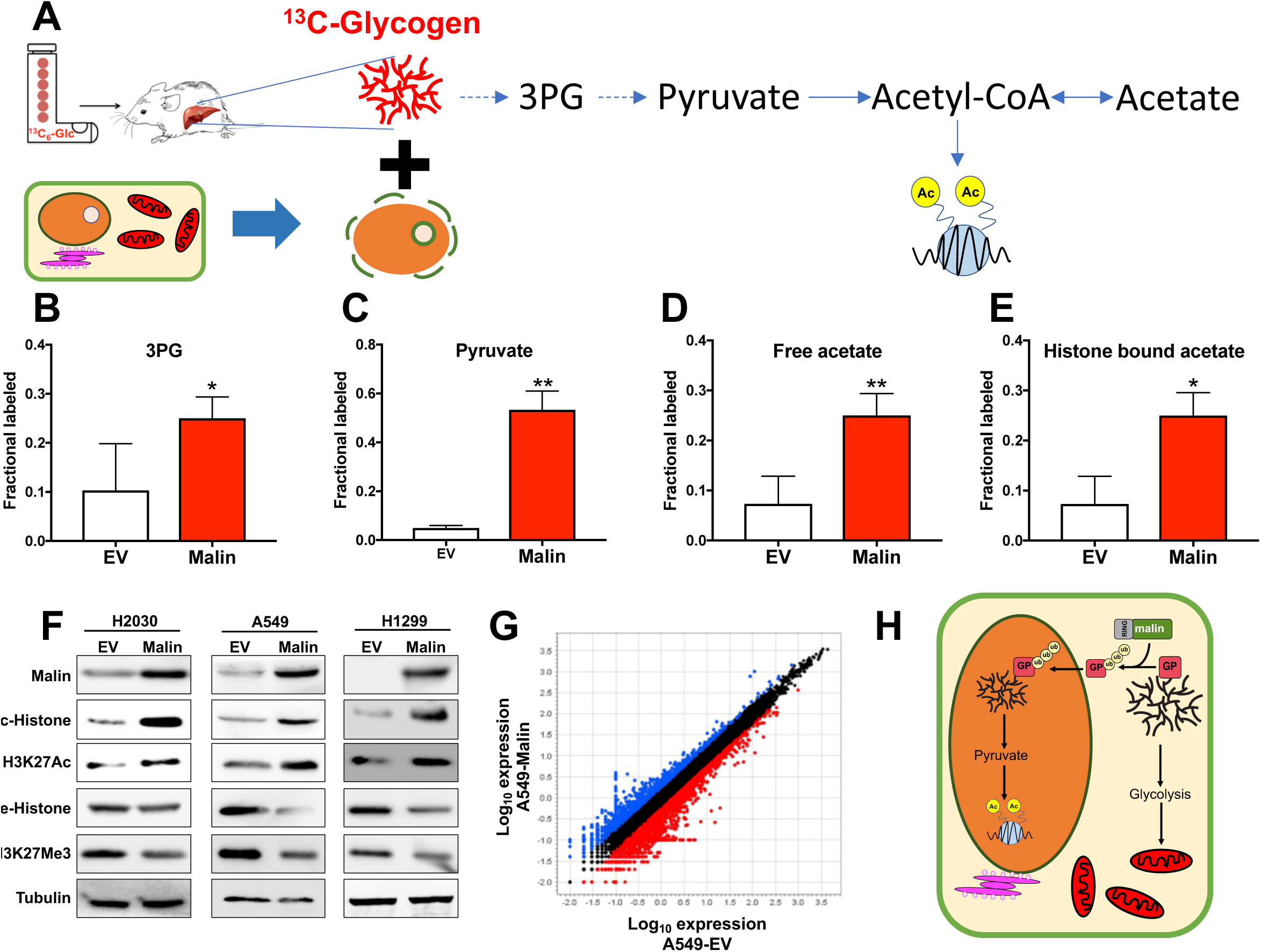
(**A**) Diagram representation of the IO-SIEM to study nuclear glycogen metabolism using ^13^C-glycogen. (**B**)-(**D**) ^13^C-enriched free 3-phosphoglycerate (3PG), pyruvate and acetate in isolated nuclei. (**E**) ^13^C-enriched histone bound acetate after histone purification followed by acid hydrolysis. (**F**) Immunoblotting analysis of histone methylation and acetylation in total histone and H3K27. (**G**) Scatter plot of the transcriptome differences between A549-empty vector (EV) and A549-Malin overexpression (Malin) by RNAseq analysis. (**H**) Diagram representation of malin regulation of nuclear glycogen metabolism and its contribution to epigenetic changes. ^*^0.01 < *P* < 0.05; ^**^ 0.001 < *P* < 0.01; ^***^ *P* < 0.001, two-tailed *t*-test (see Methods)

GP is a novel malin-interacting protein and both the liver isoform (GP_LL_) and brain isoform (GP_BB_) co-immunoprecipitated with malin (supplement Fig. 3G and H). Interestingly, malin overexpression resulted in a consistent increase in GP levels, prompting us to further investigate this interaction (Fig. 2A). The incubation of recombinant malin with ubiquitin-activating enzyme (E1), ubiquitin-conjugating enzyme (E2), ATP, and ubiquitin resulted in GP ubiquitination *in vitro* (Fig. 2B). GP_LL_ is largely localized to the cytoplasm (Fig. 2C) and co-expression with malin did not result in GP proteasomal-directed degradation, but rather translocation of GP from the cytoplasm to the nucleus (Fig. 2D and supplement Fig. 4A and B). Conversely, co-expression of a malin point mutation in either the malin RING or NHL domains abolished the GP nuclear localization (Fig. 2D). Moreover, leptomycin treatment, a nuclear exportin inhibitor^11^, resulted in the accumulation of endogenous GP in the nucleus, suggesting GP translocation is part of normal cellular physiology (supplement Fig. 4C). GP translocation to the nucleus resulted in degradation of nuclear glycogen (Fig. 2E).

**Figure 4:**
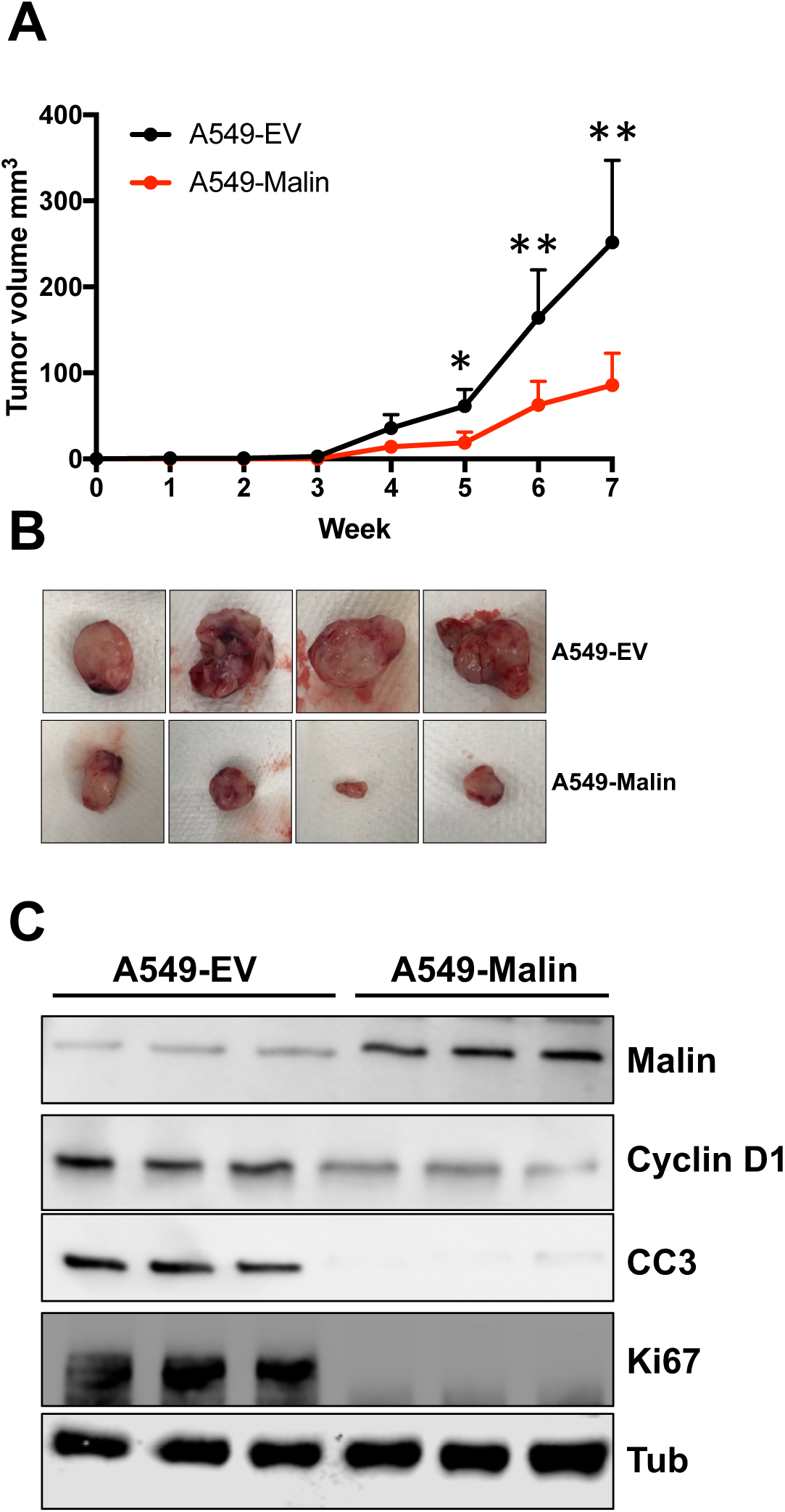
(**A**) A549 cells with malin over-expression (A549-Malin) growth is much slower than A549-empty vector (EV) cells when introduced as xenografts in immunocompromised mice. (**B**) Tumor size of A549-EV and A549-Malin xenografts at the time of sacrifice. (**C**) Immunoblotting analysis of proliferation markers Ki67, cyclin D1, and apoptotic marker cleaved caspase 3 (CC3) with tubulin (Tub) as a loading control in A549-EV and A549-Malin xenografts. ^*^0.01 < *P* < 0.05; ^**^ 0.001 < *P* < 0.01; ^***^ *P* < 0.001, two-tailed *t*-test (see Methods)

Glycolytic enzymes have been shown to moonlight in the nucleus with many having undefined functions^12,13^. Moreover, pyruvate^14,15^ and acetate^16,17^ are the primary substrates for histone acetylation, but the source for both substrates in the nucleus remains uncertain. We hypothesized that the existence of nuclear glycogen could be a separate carbon pool from the cytosolic fraction, supplying metabolic substrates for histone modification without mobilizing cytosolic metabolites. Therefore, we investigated the functional role of nuclear glycogen and GP in pyruvate and acetate production and histone modification using *In-Organello* Stable Isotope-Enriched Metabolomics (IO-SIEM) (Fig. 3A).

We generated ^13^C-glycogen using a liquid diet stable tracer enrichment method recently described^18^. 8-week old mice were fed a ^13^C_6_-glucose enriched liquid diet for 48 hours (supplement Fig. 5A) and liver glycogen was purified by trichloroacetic acid extraction and ethanol precipitation^19^. Next, the nucleus from empty vector (EV) or malin-OE cells were isolated in hypotonic cell lysis buffer. We found that pyruvate is transported into the nucleus, but neither liver glycogen nor glucose cross the nuclear membrane (supplement Fig. 5B-F). We then used a nuclear hypotonic lysis buffer to release nuclear enzymes while preserving enzyme integrity. Nuclear lysates and ^13^C-glycogen were incubated in respiration buffer for 6 hours and polar metabolites and their isotopologues were derivatized and analyzed by gas-chromatography mass spectrometry (supplemental Fig. 6). Malin-OE and nuclear localization of GP resulted in the successful breakdown of glycogen and the enrichment of glycolytic metabolites (Fig. 3B-C and supplemental Fig. 6B-E) such as 3PG and pyruvate. TCA cycle metabolites such as citrate, fumarate, succinate as well as serine and glycine were also identified in the nucleus but not enriched upon nuclear glycogenolysis (supplement Fig. 6G-M). Strikingly, both free and histone-bound ^13^C-acetate were markedly increased with malin expression (Fig. 3D and E). The enrichment of histone-bound ^13^C-acetate was confirmed by immunoblotting showing a dramatic increase in pan-histone acetylation and a corresponding decrease in pan-histone methylation in malin-OE cell lines (Fig. 3F). This epigenetic change is further validated by increased acetylation and decreased trimethylation on H3K27 (Fig. 3F). Importantly, H3K27 methylation has been implicated in lineage switching during NSCLC tumorigenesis^20,21^. These data reveal nuclear glycogen as a metabolite source for histone modification, specifically acetylation.

Epigenetic programs, i.e. chromatin organization, dictate cell fate^22^, control cell cycle^23^ and contribute to tumorigenesis^24^. Global changes in histone methylation and acetylation with malin-OE suggest a shift in the transcriptional landscape. We performed RNAseq analysis to define transcriptional changes after malin-OE and found that all three cell lines showed similar changes with an average of 1761 genes that were downregulated and 1672 genes that were upregulated more than 1.5-fold (Fig. 3G and supplement Fig. 7A and B). Principle component analysis (PCA) and clustering heat map analysis placed the EV and malin-OE lines in two separate groups (supplement Fig. 7C and D). Genes upregulated in malin-OE versus EV were enriched for pathways involved in metabolism and cell division that influence NSCLC growth. Increases in glutathione metabolism and amino acid metabolism such as arginine, tryptophan and histidine metabolism are also observed with malin-OE (supplement Fig. 7E). Downregulated genes with malin-OE were enriched for glycolysis, oxidative phosphorylation, and o-glycan biosynthesis (supplement Fig. 7E). Interestingly, genes involved in DNA demethylation and alkylation were upregulated, while genes involved in chromosome segregation, cell proliferation, and execution of apoptosis were downregulated with malin-OE (supplement Fig. 7F).

Histone hypermethylation and hypoacetylation are contributing factors to tumorigenesis^25^. Therefore, increased histone acetylation via malin-OE would potentially have anti-proliferative effects. The NSCLC cell lines A549, H1299, H2030 with malin-OE or EV were grown as xenografts in immunocompromised mice to assess tumor growth *in vivo*. A549 cells with malin-OE grew significantly slower than the EV cell line (Fig 4A-B). Similar anti-proliferative effects were seen in H1299 and H2030 cells carrying malin-OE (supplemental Fig. 8A-B). Histochemical analysis of these tumors displayed a decrease in the proliferative marker Ki67 in malin-OE cell lines and a similar decrease in the apoptosis marker cleaved caspase 3 (CC3) (Fig 4C and supplement Fig. 8C). Furthermore, we did not detect any glycogen in malin-OE tumors, confirming the contribution of malin in glycogenolysis *in vivo* (supplement Fig. 8C).

This study identifies compartmentalized, nuclear glycogenolysis as a carbon source of acetate for histone acetylation. The pathway is dependent on the E3 ubiquitin ligase malin and its ubiquitination of GP. This process represents a new concept in sub-cellular organelle communication and signaling by an E3-ligase through glycogen. Compartmentalized glycogenolysis downstream of malin-GP signaling is a crucial component of the transcriptional regulatory machinery. The inability to carry out glycogenolysis in NSCLC from lack of GP results in a lack of substrate for histone acetylation contributing to the altered epigenetic landscape seen in NSCLC.

These data also provide a context to understand emerging themes in cancer biology that connect metabolism with epigenetic regulation through undefined links. Mutations in metabolic enzymes such as isocitrate dehydrogenase, fumarate hydratase, and succinate dehydrogenase result in over-production of “oncometabolites” such as 2-hydroxyglutarate (2HG)^26^, fumarate^27^, and succinate^28^, respectively. Studies suggest that oncometabolites contribute to cancer malignancy through inhibition of histone and DNA demethylases. In addition to oncometabolites, it is known that pyruvate/acetate and serine/methionine are donors for histone acetylation and methylation, respectively. Dysregulation of acetylation and methylation contributes to tumorigenesis. While the folate pathway is well described to support methylation, the origin of compartmentalized pyruvate and acetate had remained ambiguous, especially with previous work showing nuclear and cytosolic acetyl-CoA pools are maintained separately with limited equilibration between them^16^. To date, histone hypoacetylation has been attributed to the effects of histone deacetylases (HDACs) by DNA hypermethylation^29,30^. *In organello* glycogenolysis is yet another mechanism by which cancer can suppress histone acetylation and coordinate synergistically with HDACs to drive tumorigenesis. Downregulation of malin in NSCLC drives cellular proliferation by preventing nuclear glycogen degradation for histone acetylation. This study identifies the molecular mechanism of a half-century old observation and defines the role of nuclear glycogen beyond a simple energy cache to regulating epigenetics through acetylation. Given the significant reduction in mouse xenograft growth, the glycogen metabolic pathway represents a novel therapeutic target for NSCLC.

